# Characterization of Stanniocalcin-1 Expression in Macrophage Differentiation

**DOI:** 10.1101/2020.05.18.101808

**Authors:** Cherry CT Leung, Chris KC Wong

## Abstract

Human stanniocalcin-1 (STC1) is a paracrine factor associated with inflammation and carcinogenesis. The role of STC1 in the pro- and anti-inflammatory functions of differentiating macrophage, however, is not clear. In this study, our data showed that PMA treatment induced human leukemia monocytic cells (ThP-1) differentiation to M0 macrophages. The differentiation was accompanied by a significant increase of mRNA expression levels of STC1, the pro-inflammatory cytokine TNFα, and anti-inflammatory markers, CD163 & CD206. An intermitted removal of PMA treatment reduced the mRNA levels of STC1 and TNFα but had no noticeable effects on the anti-inflammatory markers. The correlation in the expression of STC1 and pro-inflammatory markers in differentiating macrophages was investigated, using siRNA_STC1_-transfected PMA-induced cells. Consistently, the transcripts levels of TNFα and IL-6 were significantly reduced. Moreover, LPS/IFNγ-induced M1-polarization showed remarkably higher expression levels of STC1 than IL-4/IL-13-induced M2-macrophages and PMA-induced M0-macrophages. Transcriptomic analysis of siRNA_STC1_-transfected M1-polarized cells revealed an upregulation of TBC1 domain family member 3 (TBC1D3G). The gene regulates the payload of macrophage-released extracellular vesicles to mediate inflammation. The conditioned media from siRNA_STC1_-transfected M1-polarized cells were found to reduce Hep3B cell motility. The data suggest that the expression of STC1 were associated with macrophage differentiation, but preferentially to M1 polarization.

## 1. Introduction

Inflammation is one of the cancer hallmarks and is a crucial factor in the regulation of tumor progression. In tumor microenvironment (TME), cancer cells maintain intricate interactions with surrounding stroma (Poh & Ernst 2018). The infiltration of immune cells and secretion of various growth factors/cytokines modulate the process of tissue remodeling in cancers. One crucial component of tumor stroma is macrophages that multiple aspects of coordination in immune responses. Besides, macrophages can reversibly alter their endotypes to augment antitumor reactions or antagonize cytotoxic actions (Murray *et al.* 2014). Macrophages can be classified into two groups: classically activated macrophages (CAM/M1) and alternatively activated macrophages (AAM/M2). In general, M1 macrophages are considered to be pro-inflammatory and exhibit antitumor effects. On the contrary, M2 macrophages assist a pro-tumoral role by suppressing the local immune response, to promote angiogenesis, invasion, and metastasis (Biswas *et al.* 2013; Biswas & Mantovani 2010; Sica *et al.* 2008). Nonetheless, clinical and experimental data indicated that cancer tissues with high infiltration of tumor-associated macrophages were associated with poor prognosis and exhibited resistance to therapeutics (Petty & Yang 2017). An understanding of the factors responsible for macrophage homing and differentiation is recognized to be an essential immunotherapeutic strategy.

Human stanniocalcin-1 (STC1), an autocrine/paracrine factor, is suggested to play roles in inflammation and carcinogenesis. Early clinicopathological studies showed its differential expressions in paired normal and tumor tissues (Chang *et al.* 2003). Experimental characterization on the role of STC1 in some cancer hallmarks, like proliferation, apoptosis, and angiogenesis in different cancer cell models, was reported (Yeung *et al.* 2012). Later studies revealed stimulation of STC1 expression under oxidative stress in the tumor microenvironment (Prockop & Oh 2012; Yeung *et al.* 2005). The involvement of STC1 in wound healing (Yeung & Wong 2011), and cardiovascular inflammation were demonstrated (Chen *et al.* 2008). An anti-inflammatory action of STC1 was shown via the suppression of superoxide production at glomerulonephritis (Huang *et al.* 2009). The effect of STC1 on an inhibition of ROS production via mitochondrial uncoupling protein 2 in renal ischemia/reperfusion injury in mice was reported (Huang *et al.* 2012). Moreover, the counteracting effects of STC1 on LPS-induced lung injury (Tang *et al.* 2014) or ischemic cardiac injury in mice (Mohammadipoor *et al.* 2016), via inhibition of inflammatory cascades or suppression on monocytes/macrophages recruitment were revealed respectively. The anti-inflammatory effects of STC1 on other models, like osteoarthritis (Wu *et al.* 2019), and retinopathy (Dalvin *et al.* 2020) were reported.

Our previous studies showed the inhibitory effects of STC1 on the growth of STC1-overexpressing human hepatocellular carcinoma (HCC) cells in a mouse xenograft model (Yeung *et al.* 2015). The inhibitory effects of STC1 on p70^S6K^/p-rpS6 signaling in the HCC cells to reduce tumor growth was demonstrated (Leung & Wong 2018). In considering the intricate interactions at the TME, the role of STC1 in macrophage differentiation has not been addressed. With hindsight, STC1 and macrophages are known to be involved in inflammation and carcinogenesis. In this study, we hypothesized that STC1 might play a role in differentiation and inflammatory functions of macrophages and thus modulate tumorigenicity through macrophage-cancer cell interactions. Using human leukemia monocytic cell line ThP-1, we aimed to characterize the role of STC1 in macrophage differentiation and functions.

## 2. Materials and Methods

### 2.1 Cell Culture and Macrophage Differentiation

Human leukemia monocytic cell line, ThP-1 was cultured in RPMI1640, supplemented with 10% heat-inactivated fetal bovine serum and antibiotics (25 U/mL penicillin and 25 µg/mL streptomycin) (Gibco, Life Technologies). The cells were maintained in a CO_2_ (5%) incubator at 37 °C.

The cells were treated with phorbol 12-myristate 13-acetate (PMA) (Abcam) for 24 - 48 hr to stimulate macrophage differentiation (M0). An increasing concentration of PMA (2.5 – 100 nM) was applied to the cells to determine the optimal dose of the stimulation. Once the optimal PMA concentration was identified, the differentiated cells (M0) could then be further treated with selected stimulators to obtain M1 and M2 polarized macrophages. For M1 polarization, ThP-1 cells were treated with 5 nM PMA for 6 hr, followed by 20 ng/mL IFNγ (Gibco, Life Technologies), and 100 ng/mL LPS (Invitrogen; Thermo Fisher Scientific) stimulation for another 18 hr. For M2 polarization, ThP-1 cells were treated with 5 nM PMA for 6 hr, followed by 20 ng/mL IL-4, and 20 ng/mL IL-13 (Gibco, Life Technologies) stimulation for another 18 hr. The identities of M0, M1, and M2 polarized cells were characterized using both pro- and anti-inflammatory markers.

### 2.2 Total RNA Extraction and Real-time PCR

Cellular RNA was extracted by TRIZOL Reagent (Gibco/BRL) according to the manufacturer’s instruction. The A_260_/A_280_ of total RNA was >1.8, which was utilized to synthesize cDNA using SuperScript VILO Master Mix (Invitrogen, Life Technologies). Real-time PCR was performed using the Fast SYBR Green Master Mix (Applied Biosystems). The primer sequences were listed in Table 1.

### 2.3 Western Blot Analysis

Cells were lysed in a cold radio-immunoprecipitation assay (RIPA) buffer (150mM NaCl, 50mM Tris-HCL, pH 7.4, 2mM EDTA, 1% NP-40, 0.1% SDS), then were sonicated using Bioruptor Plus (Diagenode), and were centrifuged at 12,000 g for 10 min at 4 °C. Protein concentrations of the supernatant were measured by DC Protein Assay Kit II (Bio-rad) at absorbance 750 nm using a microplate reader (BioTek). Protein lysates were then resolved in SDS-PAGE and transferred onto PVDF membrane (Bio-Rad). The membrane was blocked with 5% non-fat milk in PBST for 1 hr, followed by incubation with a primary antibody and HRP-conjugated secondary antibody (Table 2). The blots were washed in PBST (3 x 15 min) after the incubation with the antibody. Specific bands were visualized using WESTSAVE Up (AbFrontier).

### 2.4 STC1 Knock-down by siRNA

On-Target human STC1 siRNA (Dharmacon) was used to knock down the STC1 transcript. Non-targeting siRNA: UGGUUUACAUGUCGACUAA, UGGUUUACAUGUUGUGUGA, UGGUUUAC AUGUUUUCUGA, UGGUUUACAUGUUUUCCUA, and human STC1 siRNA: AAACGCACAUCCCAUGAGA, GGGAAAAGCAUUCGUCAAA, GUACAGCG CUGCUAAAUUU, CAACAGAUACUAUAACAGA were used in this study. Briefly, 1×10^5^ ThP-1 cells were seeded in 24-well plates, then transfected with 50 nM siRNA using 0.5 µl Lipofectamine 2000 transfection reagent (Invitrogen; Life Technologies) in antibiotic-free complete medium. After 6 hr incubation, the siRNA transfected cells were treated with 5 nM PMA to induce M0 macrophage differentiation. In some experiments, the siRNA transfected M0 macrophages were further stimulated to obtain M1 or M2 polarized cells.

### 2.5 Boyden Chamber-based Cell Migration Assay

In siRNA transfection experiments, ThP-1 cells were treated with either siRNA_CTRL_ or siRNA_STC1_ before the stimulation of M1 or M2 polarization in quadruplicate. The conditioned media of M1 and M2 macrophages were collected, filtered, and used in the cell migration assay. The conditioned media were transferred to 24-well companion plates. The human hepatoma, HepB3 cells were seeded onto 24-well cell-culture inserts (Falcon) with a membrane pore size of 8µm in serum-free DMEM medium. After 24 hr of incubation at 37 °C, cells on the top side of the inset-membrane were removed by a cotton swab. The cells migrated to the bottom of the membrane were fixed with ice-cold methanol for 15 min, followed by 0.5 % crystal violet (Farco Chemical Supplies) at room temperature for 15 min. The membranes were then mounted. Migrated cells were captured by a microscope and counted by Image J.

### 2.6 Transcriptomic Analysis

Total RNA of siRNA_CTRL_ or siRNA_STC1_-transfected M1 macrophages was extracted. The RNA quality was measured by Agilent 2100 Bioanalyzer system. Four replicates per treatment with RNA Integrity Number (RIN) > 8 were used for library construction. DNA sequencing was conducted at the Beijing Genomics Institute (Wuhan, China) using the BGISEQ-500RS sequencer. Single-end reads of 50 bp read-length were sequenced and trimmed according to BWA’ s–q algorithm. Quality-trimmed sequence reads were mapped to human genome reference (GRCh38/hg38). Read-count data were then subjected to differential expression analysis using the edgeR package (Robinson *et al.* 2010). Genes with FDR < 0.05 were considered as differentially expressed genes (DEGs).

### 2.7 Statistical Analysis

Statistical analysis was conducted using SigmaPlot version 12.0. Data were evaluated by the Student’s t-test or one-way analysis of variance (ANOVA) followed by Duncan’s multiple range test. All data are presented as a statistical mean ± SD. A p-value < 0.05 was used as the cutoff for statistical significance.

## 3. Results

### 3.1 Dose- and time-dependent effects of PMA on ThP-1 differentiation and STC1 expression

ThP-1 monocytes were stimulated to differentiate into M0 macrophages by the treatment with an increasing dose (2.5 – 100 nM) of phorbol 12-myristate 13-acetate (PMA) for 24 – 48 hr. The cells were transformed from a suspension to an adherent form, demonstrating a differentiated phenotype (Fig. 1A). To characterize the molecular phenotypes of the differentiation, the mRNA expression levels of pro- and anti-inflammatory markers, TNFα, CD163 and CD206 were measured. Dose- and time-dependent inductions in the expression levels of the markers were noted at 24 hr (Fig. 1B) and 48 hr post-treatment (Fig. 1C). A significant increase of STC1 mRNA (Fig. 1B-C) was observed.

**Figure 1.**
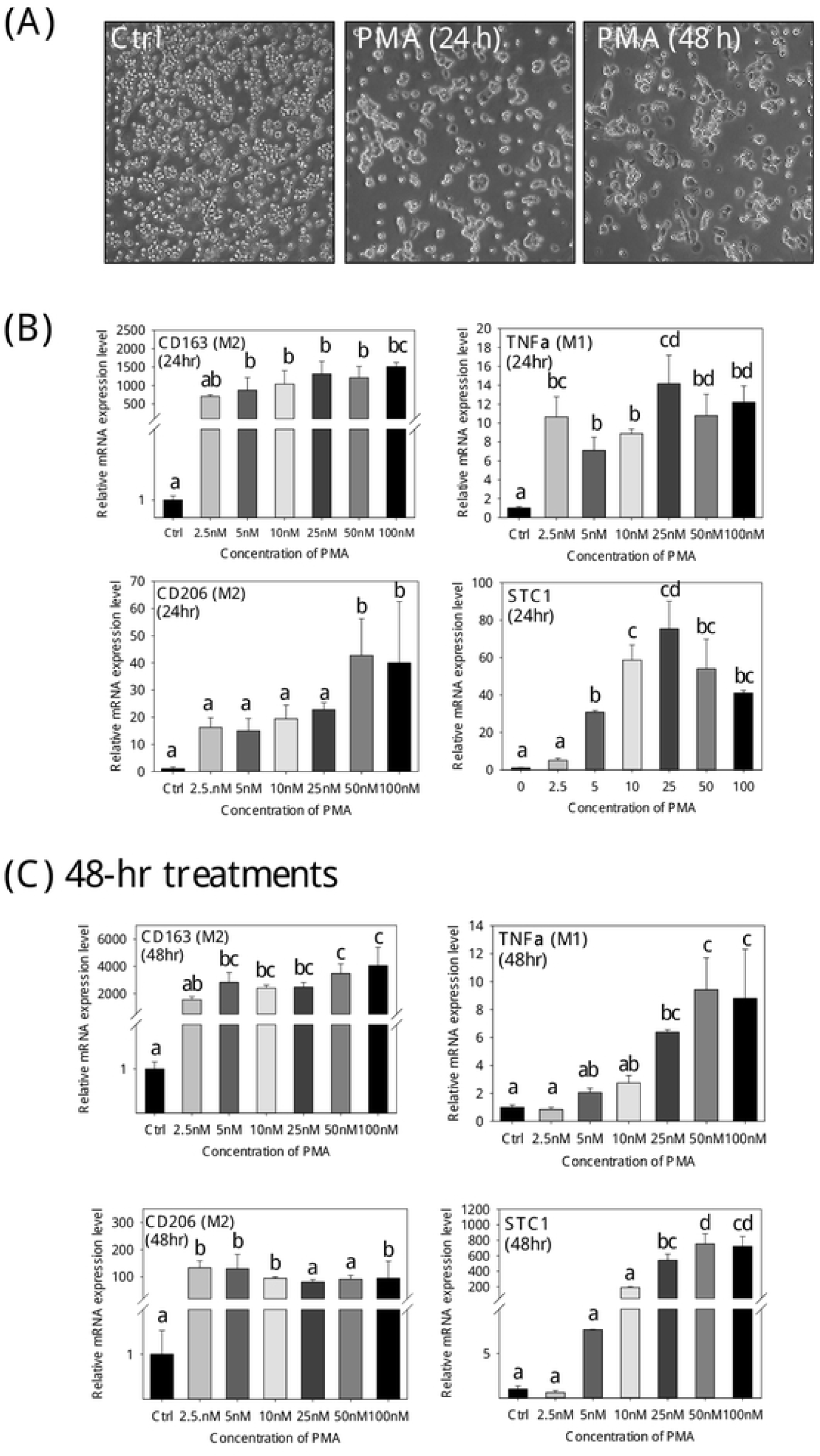
Dose- and time-dependent effects of PMA on ThP-1 differentiation to M0 macrophage and STC1 expression. **(A)** Cell morphology of ThP-1 cells changed from suspension to adherent phenotype after 5 nM PMA treatment for 24 and 48 hr. Gene expressions of CD163, TNFα, CD206, and STC1 of ThP-1 were up-regulated after **(B)** 24-hr and **(C)** 48-hr PMA treatments. Data were presented as the mean ± S.D. Bars with the same letter are not significantly different according to the results of one-way ANOVA followed by Duncan’s multiple ranges tests (p < 0.05).

An experiment with an intermitted removal of PMA in the treatment was conducted. In the cells treated with 5 nM PMA for 24 hr, followed by another 24 hr of incubation in a PMA-free medium. The cells maintained a lesser differentiated phenotype, as compared with the cells maintained in PMA-medium for 48 h, with or without washing step (Fig 2A, the right panels). The observation revealed that the intermitted removal of PMA led to a reduction of cell spreading, suggesting further differentiation was halted. Moreover, the expression levels of the pro- and anti-inflammatory markers were determined. The cells with the intermitted removal of PMA exhibited significantly lower expression levels of the pro-inflammatory cytokine TNFα and STC1 transcripts (Fig. 2B). The expression levels of the anti-inflammatory markers CD163 and CD206, however, were not noticeably affected.

**Figure 2.**
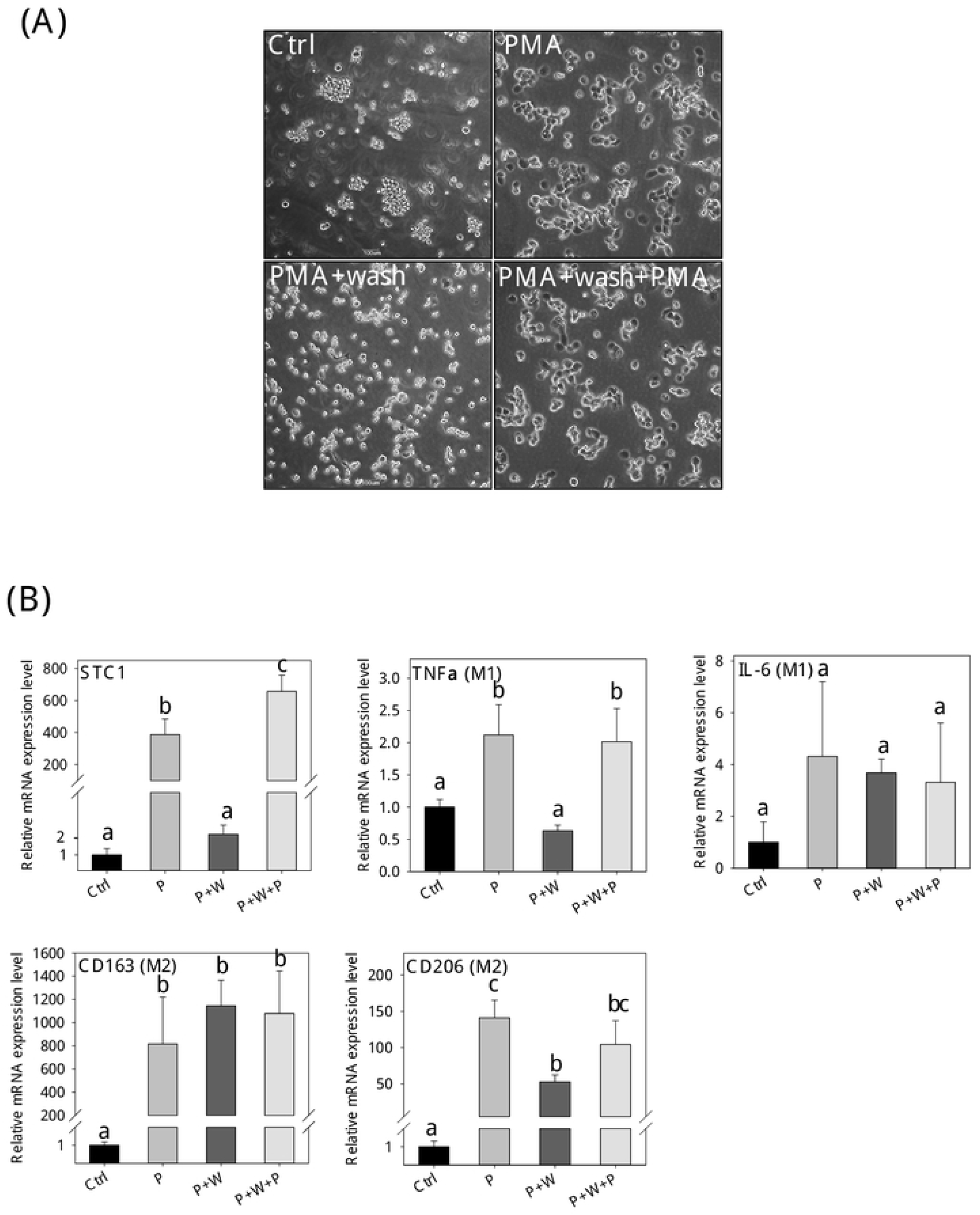
Intermitted PMA removal abrogated STC1 gene expression. **(A)** Cell morphologies of the (i) control, (ii) PMA, (iii) PMA withdrawal after 24-hr (PMA + wash), and (iv) PMA withdrawal after 24 hr + replenishment of fresh PMA (PMA + wash + PMA). **(B)** The expressions of STC1, cytokine markers (TNFα, IL6, CD163, CD206) in differentiating cells after the treatments for 48 hr (P: PMA; W: wash). Data were presented as the mean ± S.D. Bars with the same letter are not significantly different according to the results of one-way ANOVA followed by Duncan’s multiple ranges tests (p < 0.05).

### 3.2 Effects of STC1-knockdown on the expression of pro- and anti-inflammatory markers in PMA-induced differentiation (M0)

ThP-1 cells were transfected with either siRNA_CTRL_ or siRNA_STC1_ before the PMA treatment. After transfection, there were no noticeable differences in the cell phenotypes, when comparing the siRNA_CTRL_ and siRNA_STC1_ in control (Fig. 3A, the left panels) or in the PMA-treatment (Fig. 3A, the right panels). The transfection efficiency was illustrated in the siGLO co-transfection (Fig. 3B). The transfection of siRNA_STC1_ in PMA-induced ThP-1 cells (M0) showed a significant reduction in the expression levels of STC1 and the pro-inflammatory (TNFα, IL-6) and anti-inflammatory (CD163) markers (Fig. 3C).

**Figure 3.**
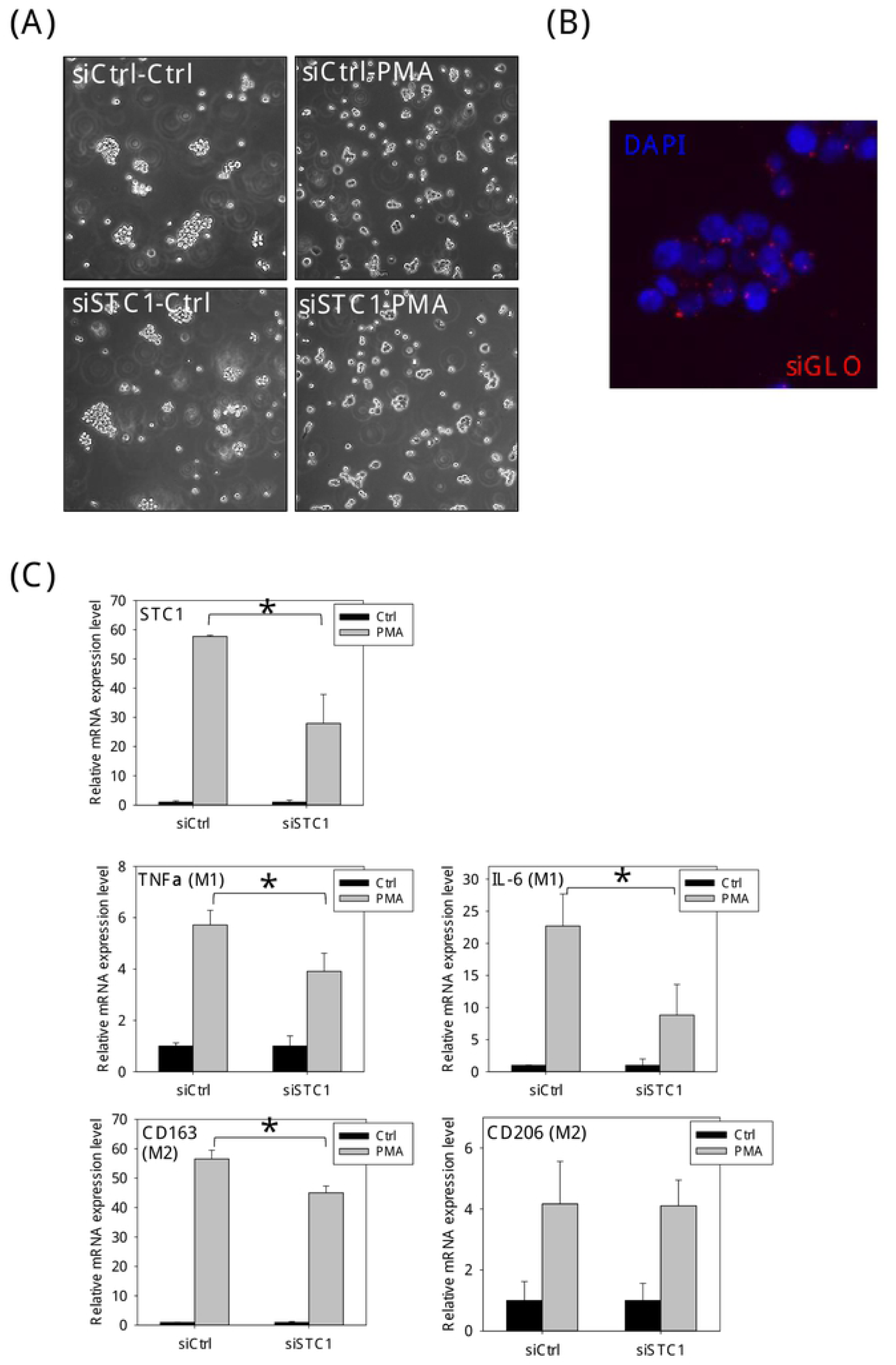
Effects of STC1-knockdown on the expression of STC1, pro- and anti-inflammatory markers in PMA-induced M0 macrophages. **(A)** Cell morphology of cells transfected with siRNA_CTRL_ or siRNA_STC1_, in the presence or absence of PMA treatment. **(B)** Transfection efficiency of siRNA was shown by siGLO (blue: DAPI; red: siGLO). **(C)** Gene expressions of STC1, cytokine markers after siRNA treatments. Significant reductions in the expression of STC1, IL-6, TNFα, and CD163 in siRNA_STC1_ transfected PMA-treated cells was observed. Data were presented as the mean ± S.D. Asterisks denote statistical significance relative to the respective control (*p < 0.05).

### 3.3 STC1 expression in M1- and M2-polarized cells

To elucidate further the association of STC1 expression in macrophage differentiation, PMA treated ThP-1 cells (M0) were further stimulated to differentiate into M1-polarized or M2-polarized cells. Figure 4A shows the cell morphology of the M0 (PMA), M1 (LPS/IFNγ), and M2 (IL-4/IL-13) polarization. M1-polarized cells exhibited epithelial-like phenotype, whereas M2-polarized cells showed adherent round-shaped morphology. The M1- and M2-polarization were characterized by measuring the expression levels of the pro- and anti-inflammatory markers. The expressions of pro-inflammatory markers (TNFα & IL-6) were remarkably induced in M1-polarized cells (TNFα: ∼150-fold; IL-6: ∼5000-fold) as compared with the levels measured in M2-polarized cells (TNFα: ∼2-fold; IL-6: ∼14-fold) (Fig. 4B). On the other hand, the expression levels of the anti-inflammatory markers (CD163: ∼400 folds; CD206: ∼8 fold) in M2-polarized cells were significantly higher, as compared with the M1-polarized cells (CD163: ∼100-fold; CD206: ∼2-fold) (Fig. 4C). In M1-polarization, STC1 gene expression was remarkably increased by 800-fold, whereas in M2-polarization, the induction was about 10-fold (Fig. 4D). Significant stimulation of STC1 mRNA expression was measured in the differentiation, in which ∼800- and∼10-fold inductions were measured in M1- and M2-polarized cells, respectively.

**Figure 4.**
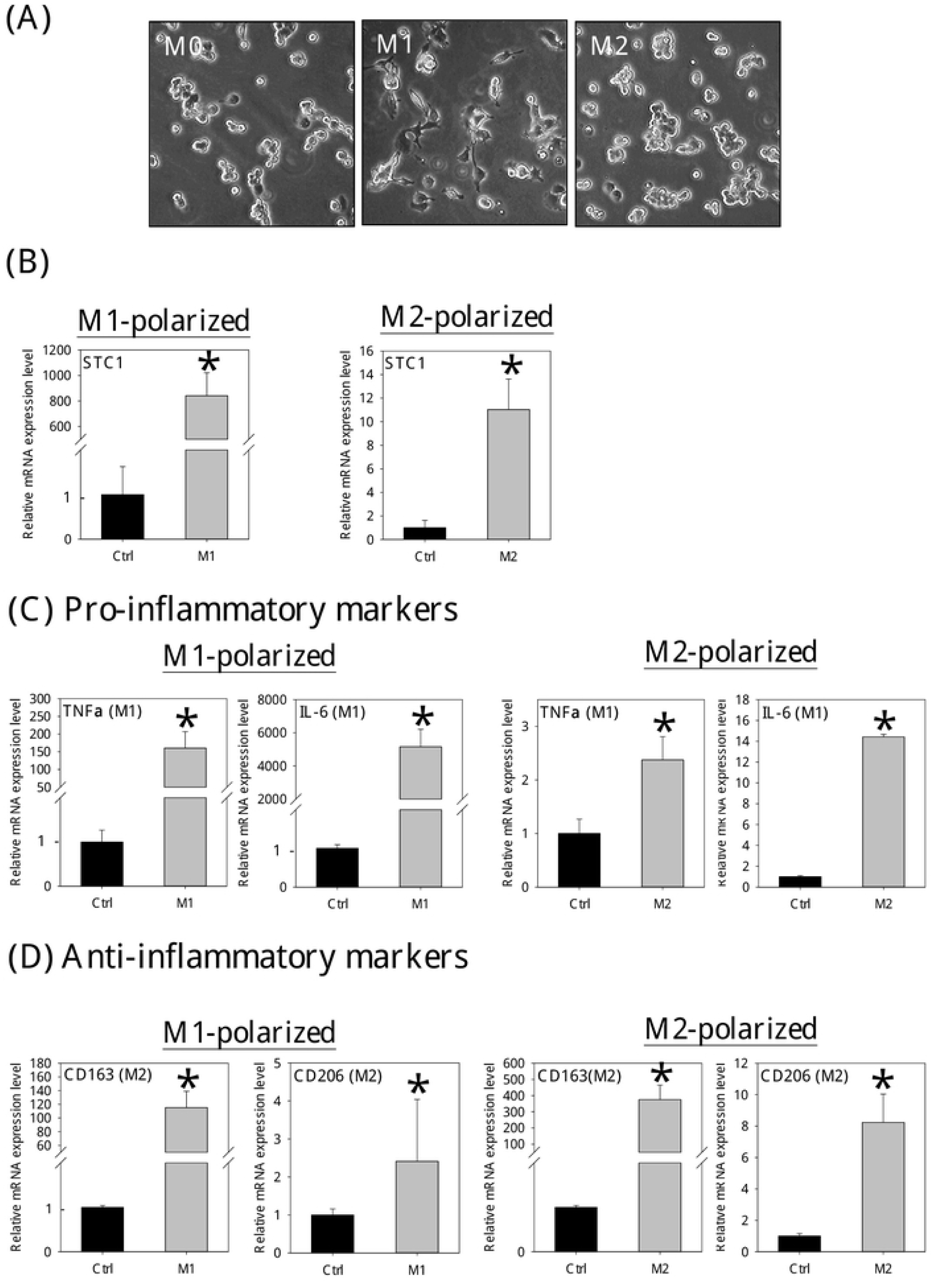
The gene expression levels of STC1 and cytokine markers in LPS/IFNγ-induced M1 or IL-4/IL-13-induced M2 macrophages. **(A)** The differentiating morphologies of M0, M1, and M2 polarized cells. **(B)** The gene expression levels of STC1 in (i) M1-polarized (the left panel) and (ii) M2-polarized (the right panel) cells. The induction of STC1 expression in M1-polarized cells was remarkably higher than in the M2 cells. Noted the different scales in the y-axes. **(C)** Messenger RNA expression levels of pro-inflammatory markers in M1- (the left panel) and M2- (the right panel) polarized cells. The induction of the pro-inflammatory markers in M1-polarized cells was remarkably higher than in the M2 cells. Noted the different scales in the y-axes. **(D)** Messenger RNA expression levels of anti-inflammatory markers in (i) M1- (the left panel) and (ii) M2- (the right panel) polarized cells. The induction of anti-inflammatory marker expression in M2-polarized cells was remarkably higher than in the M1 cells. Noted the different scales in the y-axes. Data were presented as the mean ± S.D. Asterisks denote statistical significance relative to the respective control (*p < 0.05).

### 3.4 Effects of STC1 knockdown on global gene expression in M1-polarized cells

In the transcriptomic analysis of STC1-knockdown, quality-trimmed Illumina reads of about 26.3 M were acquired from each sample of siRNA_CTRL_ and siRNA_STC1_-transfected M1-polarized cells. A volcano plot showed that only two genes were significantly dysregulated (Fig. 5A). The STC1 transcript was downregulated, while the TBC1D3G mRNA was up-regulated (Fig. 5B, the left panel). Real-time PCR analysis validated the changes of STC1 and TBC1D3G mRNA expression (Fig. 5B, the right panel).

**Figure 5.**
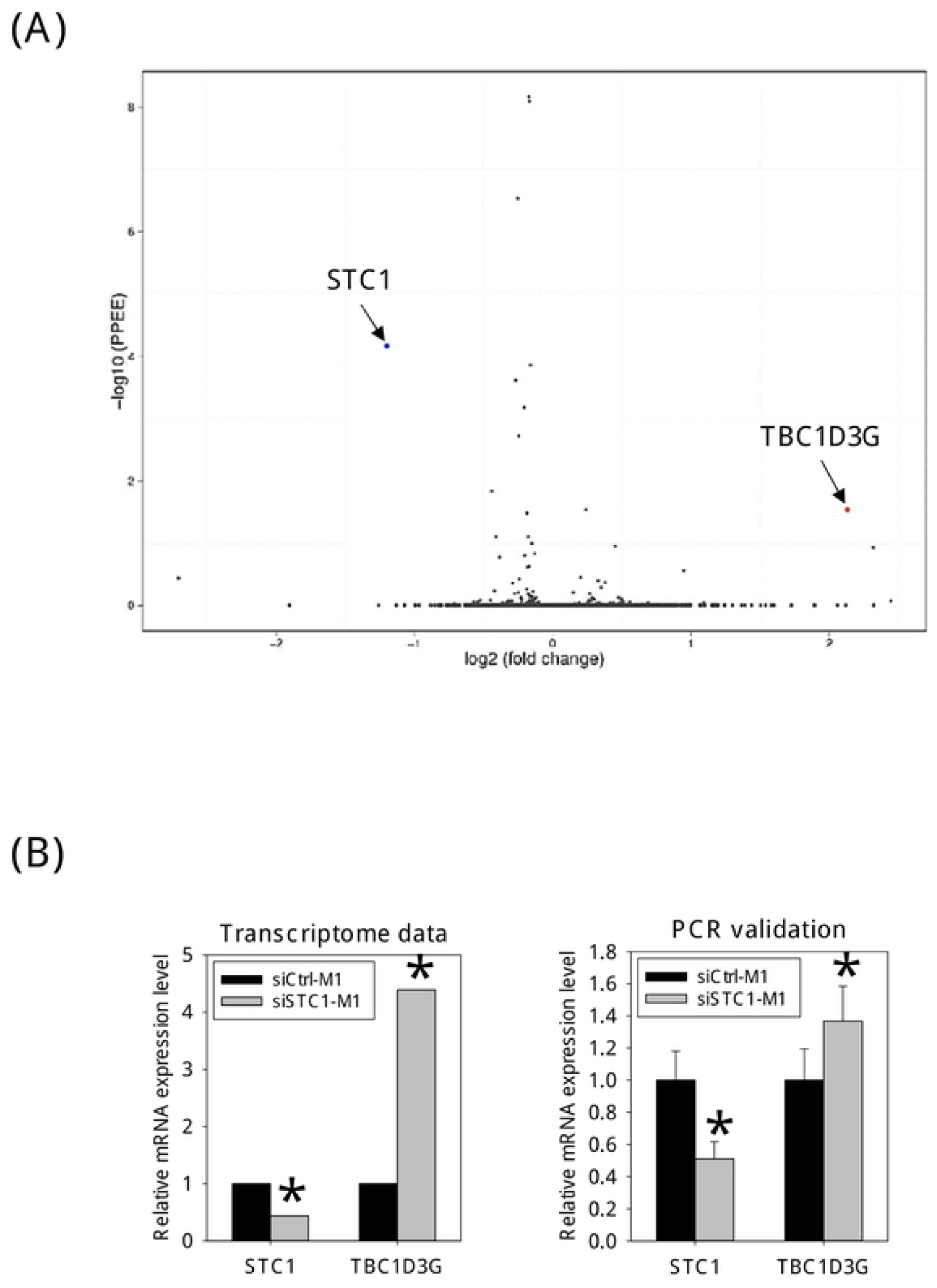
Transcriptomic analysis of STC1 knockdown M1 cells. **(A)** Volcano plot of dysregulated genes. The x-axis represents the differential gene expression (log2). Y-axis represents the p-value (-log10). A red dot represents the significantly up-regulated DEG - TBC1D3G. A blue dot represents the significantly down-regulated DEG - STC1. Gray points represent non-DEGs. **(B)** Messenger RNA levels of STC1 and TBC1D3G, as revealed by transcriptome (the left panel) and validated by real-PCR (the right panel). Data were presented as the mean ± S.D. Asterisks denote statistical significance relative to the respective control (*p < 0.05).

### 3.5 Effects of conditioned media from siRNA_STC1_ transfected M1-polarized cells on Hep3B migration

Conditioned media from the siRNA_CTRL_- and siRNA_STC1_-transfected M1-polarized cells were collected. The efficiency of siRNA delivery was checked by siGLO transfection, and the reduction of STC1 expression was measured using real-time PCR (data not shown). The conditioned media of the M1-polarized cells (siRNA_STC1_-transfected) showed inhibitory effects on the migration of Hep3B cells (Fig. 6A). On the other hand, conditioned media from STC1 knockdown M2-polarized cells showed no noticeable effects on the migration (Fig. 6B).

**Figure 6.**
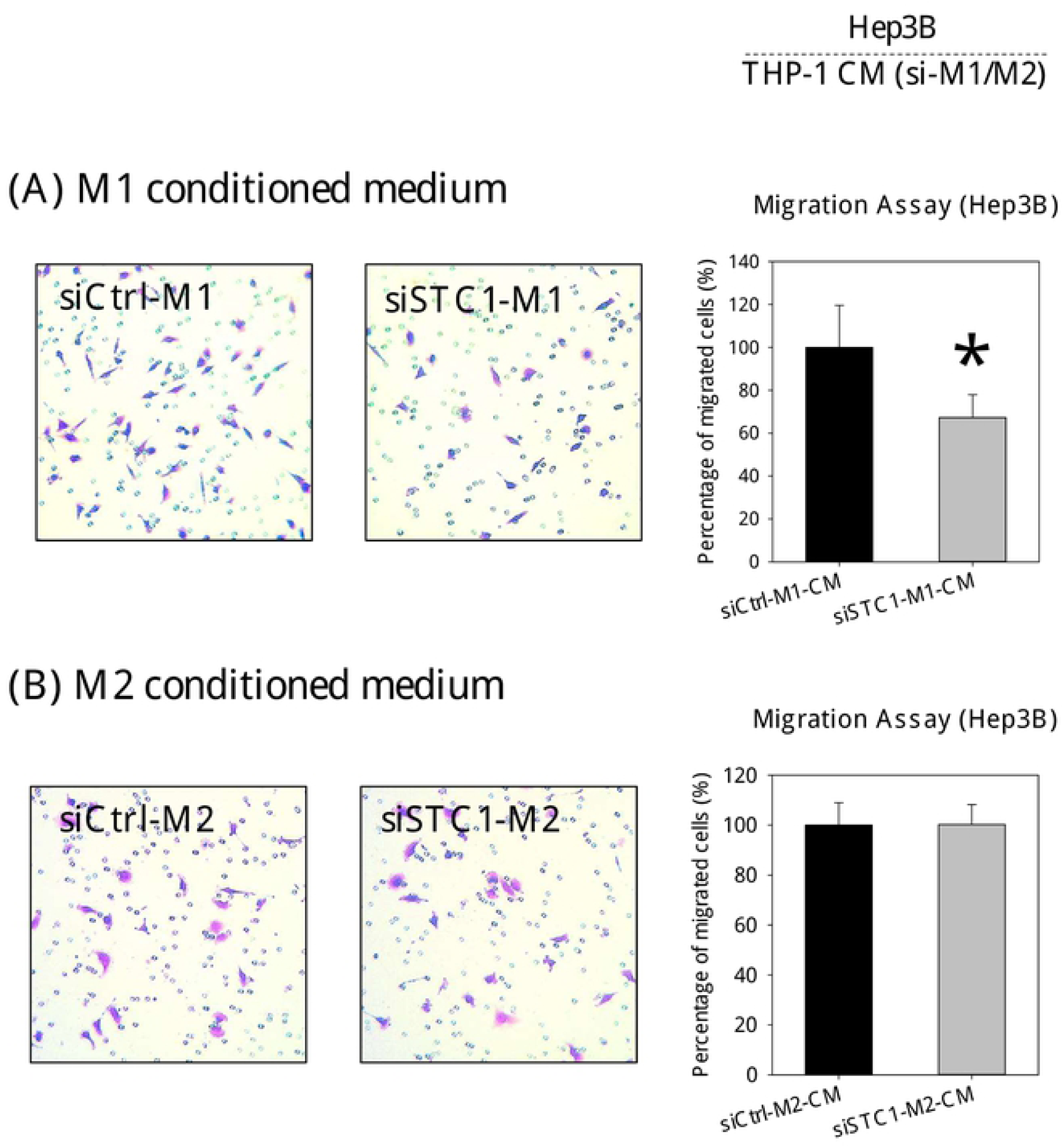
Cell Migration assay. Effects of conditioned media from siRNA_STC1_ transfected **(A)** M1-polarized cells and **(B)** M2-polarized cells on Hep3B migration. A significant reduction in the motility of STC1-knockdown M1 cells was noted. Data were presented as the mean ± S.D. Asterisks denote statistical significance relative to the respective control (*p < 0.05).

## 4. Discussion

Current evidence suggests the involvement of STC1 in inflammation, based on its inhibitory effects on the production of pro-inflammatory cytokines and chemokines, and suppressive effects on macrophage responses to chemoattractant (Sheikh-Hamad 2010; Yeung *et al.* 2012). In this study, the connection of STC1 to macrophage differentiation, through studying its association with M0-, M1- and M2-polarization was revealed. The effect of STC1-knockdown in macrophages on transwell migration of a hepatocellular carcinoma cell-line Hep3B was studied.

PMA treatment on ThP-1 cells stimulated the expression of STC1 and inflammatory markers. PMA is a protein kinase C (PKC) activator, a standard chemical used to induce differentiation of monocytic ThP-1 into macrophages via PKC activation (Lund *et al.* 2016). Multiple downstream signaling pathways were reported to be responsible for PMA-induced leukemic differentiation. Those include phosphatidylinositol 3-kinase/Ca^2+^, Raf-1, MAPK, and NF-κB signaling pathways and superoxide production (Kang *et al.* 1996; Park *et al.* 2002). The Ca^2+^-signaling and stress-induced pathways were also demonstrated to stimulate STC1 expression in various studies (Huang *et al.* 2009; Law *et al.* 2008; Nguyen *et al.* 2009; Yeung *et al.* 2003; Yeung *et al.* 2005). With hindsight, STC1 expression is known to be induced in cell differentiation. These include neural cells (Yeung *et al.* 2003; Zhang *et al.* 1998), hematopoietic cells (Serlachius *et al.* 2002; Serlachius *et al.* 2004), osteoblast cells (Yoshiko *et al.* 2003), ovarian cells (Luo *et al.* 2004) and adipocyte (Serlachius & Andersson 2004). Therefore, the observation of increased STC1 expression in macrophage differentiation is consistent. Macrophage activation comprises the resting (M0), and polarizing M1 and M2 states. The classical M1 and alternative M2 macrophages that confer pro- or anti-inflammatory activities. Our following question is on the connection of STC1 to these heterogeneous populations of macrophages.

Our data show that differentiation of ThP-1 to M0, M1, and M2 macrophages induced STC1 expression, which was considerably higher in M1 polarized cells. In examining the effects of intermitted PMA-withdrawal, macrophage differentiation (M0) was arrested. STC1 expression was significantly lower as compared with the continuous PMA-treated cells. Our observation was consistent with the report by Spano and co-workers, which showed that PMA-withdrawal caused cell de-differentiation (Spano *et al.* 2013). In the measurement of cytokine markers in the intermitted PMA treated cells, a significant reduction in the expression of the pro-inflammatory marker, TNFα were noted. In the STC1 knockdown experiment, the transcript levels of pro-inflammatory markers (TNFα, IL-6) were downregulated. The data suggested a possible correlation in the expression levels of STC1 and pro-inflammatory markers. To further elucidate this correlation, we adopt the standard protocols to stimulate M1- and M2-polarization. The identity of the differentiated macrophages was characterized using the standard markers (Orecchioni *et al.* 2019). Our data show that a remarkably higher expression of STC1 was measured in M1 polarization, as compared with M2. M1 macrophages are known to be pro-inflammatory to produce cytokines to exert tumoricidal activities (Poh & Ernst 2018). The biological function of STC1, however, was suggested to be anti-inflammatory (Sheikh-Hamad 2010). The pro-inflammatory role of M1-polarized cells and the reported function of STC1 seemed to be inconsistent. However, in considering a possible negative feedback circuity in controlling pro-inflammatory signals, the expression of endogenous anti-inflammatory mediators (i.e., STC1) may help to minimize the persistent activation of immune cells in inflammatory responses. In fact, the negative feedback regulation of inflammation has been reported for over 20 years (Hanada & Yoshimura 2002). For instance, the negative feedback regulation was reported in various inflammatory pathways, like NKκB, Smad, and JAK/STAT by the suppressors of cytokine signaling or cytokine-inducible SH2 protein (Yoshimura *et al.* 2003; Yoshimura *et al.* 2005). In our previous study in characterizing the effects of STC1 on tumorigenicity of hepatocellular carcinoma Hep3B cells, IL-6 and IL-8 treatment induced STC1 expression. However, STC1 exhibited inhibitory action on the pro-migratory functions of IL-6 and IL8 (Yeung *et al.* 2015). Other examples of reciprocal regulation in inflammation responses include glucocorticoid-cytokine (Newton *et al.* 2017) and inducible NO synthase-connexin 43 (Li *et al.* 2011). Retrospectively, this suggests that the highest expression level of STC1 in M1-polarized cells may serve as a negative mediator in inflammation.

Transcriptomic analysis of STC1-knockdown M1-polarized cells identified a deregulated gene, Tre-2/Bub2/Cdc16 (TBC1) domain family member 3 (TBC1D3). The gene is reported to regulate the payload (i.e., RNA-binding protein, secretory pathways regulatory proteins) of released extracellular vesicles (EVs) from macrophages to mediate inflammation and tissue repair (Qin *et al.* 2019). The high expression of TBC1D3 was associated with decreased STAT3 phosphorylation in recipient cells (Qin *et al.* 2019). A decreased phosphorylated level of STAT3 was found to inhibit the proliferation and migration of HCC (Xie *et al.* 2018). Consistently in this study, the incubation of Hep3B cells in conditioned media of siRNA_STC1_-transfected M1-polarized cells showed a significant reduction in cell motility. Alternatively, the high expression level of STC1 in M1-polarized cells might have low expression of TBC1D3, resulting in higher phosphorylation of STAT3 in cancer cells, which associated with poor prognosis in HCC (Chan *et al.* 2017; Yeung *et al.* 2015).

In summary, we unravel the connection of STC1 expression in macrophage differentiation, and its association with the expression of pro-inflammatory markers. STC1 may serve as a negative mediator to regulate inflammation, which might be associated with the expression of TBC1D3. The data support further investigation, using clinical HCC samples to unravel the intricate association of STC1, macrophages, and tumor cells in carcinogenesis.

## Declaration of interest

The authors declare they have no actual or potential competing financial interests.

## Acknowledgments

This work was supported by the General Research Fund [HKBU12103817], Research Grant Council, Hong Kong.

## Author contribution statement

CKCW was responsible for the design and planning of the experiments. CCTL was responsible for experimental works. Both CKCW and CCTL are responsible for data analysis and manuscript writing.

